# Development of a sequence-based *in silico* OspA typing method for *Borrelia burgdorferi* sensu lato

**DOI:** 10.1101/2023.11.30.568280

**Authors:** Jonathan T. Lee, Zhenghui Li, Lorna D. Nunez, Daniel Katze, B. Scott Perrin, Varun Raghuraman, Urvi Rajyaguru, Katrina E. Llamera, Lubomira Andrew, Annaliesa S. Anderson, Joppe W. Hovius, Paul A. Liberator, Raphael Simon, Li Hao

## Abstract

Lyme disease (LD), caused by spirochete bacteria of the genus *Borrelia burgdorferi* sensu lato, remains the most common vector-borne disease in the northern hemisphere. *Borrelia* outer surface protein A (OspA) is an integral surface protein expressed during the tick cycle, and a validated vaccine target. There are at least 20 recognized *Borrelia* genospecies, that vary in OspA serotype. Traditional serotyping of *Borrelia* isolates using OspA-specific monoclonal antibodies is technically challenging and reagent-constrained. This study presents a new *in silico* sequence-based method for OspA typing using next-generation sequence data. Using a compiled database of over 400 *Borrelia* genomes encompassing all major genospecies, we characterized OspA diversity in a manner that can accommodate existing and new OspA types and then defined boundaries for classification and assignment of OspA types based on the sequence similarity. To accommodate potential novel OspA types, we have developed a new nomenclature: OspA *in silico* type (IST). Beyond the ISTs which corresponded to existing OspA serotypes (ST1-8), we identified nine additional ISTs which cover new OspA variants in *B. bavariensis* (IST9-10), *B. garinii* (IST11-12), and other *Borrelia* genospecies (IST13-17). Compared to traditional OspA serotyping methods, this new computational pipeline provides a more comprehensive and broadly applicable approach for characterization of OspA type and *Borrelia* genospecies to support vaccine development.

**Impact Statement:** As the incidence of LD continues to rise, so does the need to maintain genomic surveillance of disease-causing *Borrelia spp.* and support clinical development of new vaccines. Towards this goal, introducing the OspA *in silico* type (IST) nomenclature scheme, as well as the open-source release of this OspA analysis pipeline, will enable characterization of novel *Borrelia* OspA types using NGS data without the need for traditional, antibody-based serotyping systems.

## INTRODUCTION

Lyme borreliosis, or Lyme disease (LD), is the most common tickborne disease in the northern hemisphere and is caused by genospecies members of *Borrelia burgdorferi* sensu lato (s.l.) (1–3). *B. burgdorferi* spirochetes are extracellular pathogens whose lifecycle is restricted to cycling between vertebrae reservoir hosts and its vector, *Ixodes* ticks (4). At least 20 accepted genospecies have been described within the species complex of *B. burgdorferi* s.l. (5, 6). Most LD cases in North America are caused by *B. burgdorferi* sensu stricto (hereafter *B. burgdorferi*), while multiple species of *Borrelia* cause the majority of LD in Europe and Asia including *B. burgdorferi B. garinii*, *B. bavariensis*, and *B. afzelii* (1–3). The number of LD cases attributable to these species has steadily increased as the disease continues to geographically expand (7–12).

Outer surface protein A (OspA), encoded by *ospA* on linear plasmid 54 (lp54), is a dominant outer membrane antigen found in all *B. burgdorferi* s.l. species. OspA is expressed by the spirochetes while in the tick mid-gut where it is integral to persistent colonization (13) and is a proven antigen target for LD vaccines (14–20). The dominant disease-causing species of *Borrelia* prevalent in North America and Europe primarily belong to six OspA serotypes (ST1-6) (21). *B. burgdorferi*, that is the dominant genospecies in the US, shows homogeneity for OspA serotype 1 (ST1). By comparison, in Europe, isolates comprise a diverse range of genomic species and associated OspA serotypes. A comparison of *ospA* sequences revealed that certain European *Borrelia* species were rather homogeneous (*e.g.*, *B. afzelii* ST2 and *B. bavariensis* ST4) while others were more heterogeneous (*e.g.*, *B. garinii* ST3, ST5, ST6) in their OspA grouping (22, 23), and that there was general agreement between *ospA* genotype and OspA serotype assigned using type-specific monoclonal antibodies (mAbs) (24).

The first seven OspA serotypes (ST1-7) reported in 1993 (24), together with ST8 in 1996 (25), were identified using a traditional serotyping system limited by the non-commercial availability of reference mAbs and a requirement for mAb combinations for identification. These limitations have led researchers to move away from serological typing methods in favor of alternative approaches (26, 27). Multilocus sequence typing (MLST) is a current gold standard genotyping tool (28), as well as typing based on the 5S–23S rDNA (*rrfA*-*rrlB*) intergenic spacer (IGS) (29). These approaches have the advantage of being more discriminative, but none have allowed reliable differentiation of the *B. garinii*-associated OspA types in Europe. Moreover, these prior approaches are not amenable to describing the diversity of OspA as it pertains to coverage by investigational OspA vaccines in development. Increasingly, classic typing approaches are being displaced by whole genome sequencing (WGS) with the rapid advancement in next-generation sequencing (NGS) platforms and sequence analysis algorithms (30). *In silico* strain typing based on WGS/NGS data can provide more precise information on the diversity of *B. burgdorferi* s.l. and the relationship between phylogenetic clusters of OspA variants, *Borrelia* genospecies, and the OspA type.

To support *Borrelia* surveillance and typing, we have compiled a database of over 400 genomes representing 11 genospecies of *B. burgdorferi* s.l.. Approximately half were accessed from PubMLST and GenBank (31) and the remainder from NGS of isolates determined in-house. The isolates in this collective database comprise human pathogenic *Borrelia* species of North America and Europe (*B. burgdorferi*, *B. afzelii, B. garinii*, *B. bavariensi*s, *B. spielmanii*, *B. valaisiana,* and *B. mayonii*) as well as those not recognized as human pathogens (*B. japonica, B. turdi, and B. finlandensis*). Our analysis of these genomes revealed a comprehensive picture of OspA diversity across and within the major pathogenic *B. burgdorferi* genospecies and has been used to define sequence similarity boundaries between OspA types. The development of a sequence-based OspA *in silico* typing (IST) scheme, described here, provides a valuable tool for characterization of clinical samples at the level of OspA type and genospecies.

## METHODS

### Sources of *Borrelia* isolate collections

*Borrelia* genomes used for this study (n = 421) were sourced from an internal collection of unique isolates (n = 193) (Table S1, PRJNA1041728) and public genome sequences (n = 228) from the PubMLST *Borrelia* isolates database (https://pubmlst.org) and GenBank (Table S2). Accession numbers of externally sourced isolates are included in Table S2.

### Bacterial growth and whole genome sequencing

MKP (Modified Kelly-Pettenkoffer) media prepared in-house (32) was used for cultivation of *B. afzelii, B. garinii, B. bavariensis* and *B. spielmanii* isolates from frozen stock vials, whereas BSK-H (modified Barbour-Stonenner Kelly) media (Sigma-Aldrich Cat# B8219) was used for the culture of *B. burgdorferi.* Cultures were incubated at 34° C for 5 to 14 days and closely monitored for bacterial growth by dark field microscopy. Cultures were centrifuged at 10,000 x g for 10 minutes once the spirochete concentration reached at least 1.0 × 10^6^ cfu/mL. Cell pellets were processed for genomic DNA extraction following the magnetic bead-based Genfind V3 DNA isolation protocol (Beckman Coulter Life Sciences Cat# C34881).Next-generation sequencing was performed on the Illumina MiSeq platform, with 2×300bp paired-end NGS chemistry following a slightly modified protocol previously described by Jones *et al.* (33). Read quality was verified using samtools (v1.15.1) to find the per-base-coverage of the chromosome. An average sequencing depth of 193x was achieved across all isolates.

### Genotype characterization and phylogenetic analysis

For NGS reads from each isolate, *de novo* genome assembly was performed using the CLC Genomic Workbench (v21.0.5) with default settings. Whole-genome core alignment was performed using Parsnp (34), and the OspA-specific phylogenetic tree was generated using MEGA11 (35). Consensus OspA sequences per IST were calculated using CLC Genomic Workbench. For ISTs where ≤ 2 sequences were identified, a single OspA variant was used in place of a consensus sequence.

### Development of OspA *in silico* typing (IST) pipeline

#### OspA reference variants and IST

Unique OspA protein sequences (Table S3) were obtained from a combination of sources: NCBI, the PubMLST database (pubmlst.org) and internal collections of *Borrelia* (Table S1, Table S2). Phylogenetic analysis (see previous section) identified distinct OspA clusters, defined here as OspA ISTs wherever there are more than 2 isolates.

#### Read alignment and consensus ospA sequence

*Borrelia* NGS reads were aligned by bwa mem (v0.7.17) to reference sequences for multiple genospecies and ISTs: *B. burgdorferi* (IST1), *B. afzelii* (IST2), *B. garinii* (IST3, IST5, IST6, IST11), and *B. bavariensis* (IST4, IST9, IST10. Per-base coverage was extracted using samtools and the mean and standard deviation coverage was computed for each alignment. The alignment with the highest mean minus one standard deviation was used for further analysis. The selected alignments were assessed for median coverage and sequence depth at each nucleotide in the *ospA* gene. Samples with under 100 reads aligned, median coverage <10X, and/or sequence depth below 5X across <99% of the gene were deemed low quality and excluded from further IST analysis. Variant calling was performed using bcftools (v1.17) with the setting QUAL>=30, and the consensus *ospA* sequence generated by bcftools.

To predict OspA IST from assembled genomic contigs, BLASTN was instead used to obtain alignments to the IST-specific reference sequences. The alignment with the highest E-value was used for downstream processing with sequences with less than 99% per-base alignment considered low quality and excluded from further analysis.

#### OspA IST assignment and estimation of IST-specific pairwise identity threshold

The consensus *ospA* nucleotide sequence was translated to amino acids and compared against 79 reference OspA variants (Table S3) using MAFFT (v.7.480 (36)). A pairwise sequence identity was then calculated for each comparison. In cases of exact sequence match, the OspA IST of the known sequence variant was used for IST assignment. For non-exact matches, the IST of the OspA variant with the highest pairwise identity was assigned only if the pairwise identity met the corresponding threshold. Otherwise, an “Unknown IST” designation was assigned.

To estimate IST-specific pairwise identity threshold, all known OspA variants were multi-aligned using MAFFT and the number of mismatches calculated using a custom Python script to determine pairwise sequence identity. For each OspA variant, the maximum pairwise sequence identity within and outside of their corresponding IST was calculated. A similarity threshold for each IST was then determined as the midpoint between the lowest within-IST percent similarity and highest out-of-IST percent similarity.

#### Evaluation of OspA NGS pipeline using clinical isolates

To evaluate the performance of the OspA typing pipeline, 22 clinical *B. burgdorferi* s.l. isolates obtained from Valneva SE (Saint-Herblain, France) were used for testing. Bacterial genomic DNA was prepared using the Illumina Nextera XT V2 kit and sequenced on the Illumina MiSeq platform.. Sequence data were processed using the *in silico* OspA typing pipeline. Results were compared with OspA serotypes previously determined by amino acid (AA) sequence alignment to reference strains containing candidate OspA sequences (20).

### Data Availability Statement

Raw data presented in this study will be deposited at NCBI SRA under accession PRJNA1041728 pending the acceptance of the manuscript. Accession numbers will be included in Table S1, and the code for *in silico* OspA typing pipeline will be available on GitHub (https://github.com/pfizer-opensource) following acceptance of the manuscript.

## RESULTS

### *Borrelia* genome collections

We sequenced and assembled the genome of 193 *Borrelia* isolates (Table S1). The majority of isolates were collected from humans between 1988 and 2018, 81% from human skin biopsy samples (Figure 1A). A small percentage (7%) of isolates were obtained directly from ticks. Overall, *B. afzelii* and *B. burgdorferi* predominated and all *B. afzelii* included in the collection were isolated from Europe (88.4% of these from the Netherlands) (Table S1). Within *B. burgdorferi*, 77.5% were collected from the US, whereas the remaining isolates were from European countries.

**Figure 1.**
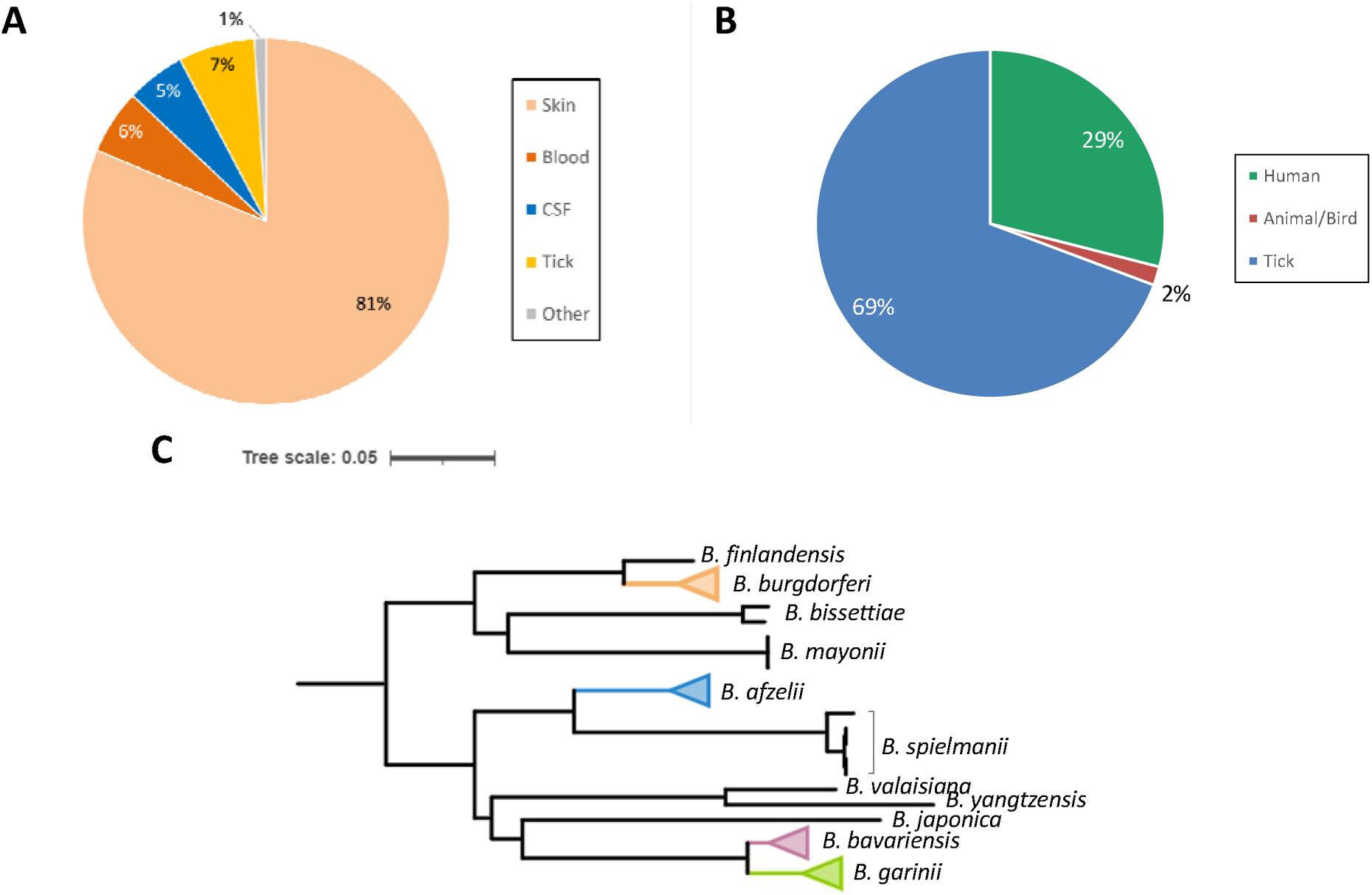
Host and genospecies distribution of *Borrelia* strains. (A) Isolates sequenced for this study were largely collected from human hosts, with the majority coming from skin biopsies. A small subset of strains was isolated directly from ticks. (B) Genomes sourced from public collections were predominantly from ticks rather than human hosts. Some isolates were obtained from non-human animal hosts. (C) Core genome phylogeny of *Borrelia* strains and representation of the predominant genospecies in the combined collection.

An additional 228 genomes were sourced from the public databases PubMLST and GenBank (Table S2). In contrast to our own sequenced collection, these were predominantly obtained directly from ticks (Figure 1B). Nearly half of the isolates in this subset were *B. burgdorferi* from North America, with the remainder predominantly consisting of *B. garinii* and *B. bavariensis* from Europe and Japan (Table S2). When combined with our internal collection, a total of 11 different *B. burgdorferi* s.l. genospecies were represented in the aggregate collection, with *B. afzelii, B. burgdorferi, B. garinii, and B. bavariensis* all well represented (Figure 1C).

### Genetic diversity of OspA within and across *Borrelia* genospecies

From the compiled genome collection, we identified a total of 79 unique OspA protein variants (Table S3). The phylogenetic tree of OspA variants was annotated to identify the existing reference *Borrelia* strains with known OspA serotypes (ST1-8) (Figure 2). As expected, *ospA* sequences sharing the same OspA serotype were phylogenetically clustered. We also observed additional clusters in the phylogeny that did not correspond to any previously reported serotypes. To accommodate these potential novel OspA types, and to better characterize the OspA diversity, we have developed a new nomenclature: OspA *in silico* type (IST). We found that OspA IST correlated strongly with OspA ST1-8 and could be linked to the corresponding genospecies: *B. burgdorferi* (IST1), *B. afzelii* (IST2), *B. bavariensis* (IST4), and *B. garinii* (IST3, IST5, IST6, IST7 and IST8) (Figure 2, Table 1). OspA variants identified from *B. burgdorferi* (IST1) and *B. afzelii* (IST2) were limited to a single phylogenetic cluster. In contrast, two other genospecies typically restricted to LD cases in Europe and Asia, *B. garinii* and *B. bavariensis*, were more heterogenous with multiple clusters of OspA variants (Figure 2). OspA variants identified among *B. bavariensis* strains were clustered into three phylogenetic groups, one of which corresponded to IST4 (sequence variant 12, *i.e.*, European OspA ST4). OspA variants corresponding to two new phylogenetic groups designated IST9 and IST10 were detected in *B. bavariensis* strains from Asia. Only OspA variant 12 was represented in the phylogenetic IST4 cluster, although 23 isolates carried this OspA variant (Table S1 and S2). The diversity of OspA variants from *B. garinii* was more considerable spanning seven phylogenetic clusters. Five corresponded to the OspA serotypes originally identified for *B. garinii* (ST3, ST5-8) (24, 25); two novel *B. garinii* clusters were labeled as IST11 and IST12. Unlike the other *B. garinii* ISTs, only a single OspA variant was identified from IST8. Finally, individual phylogenetic groups were identified for the genospecies *B. spielmanii* (IST13)*, B. mayonii* (IST14)*, B. valaisiana* (IST15)*, B. turdi* (IST16), and *B. yangtzensis* (IST17).

**Figure 2.**
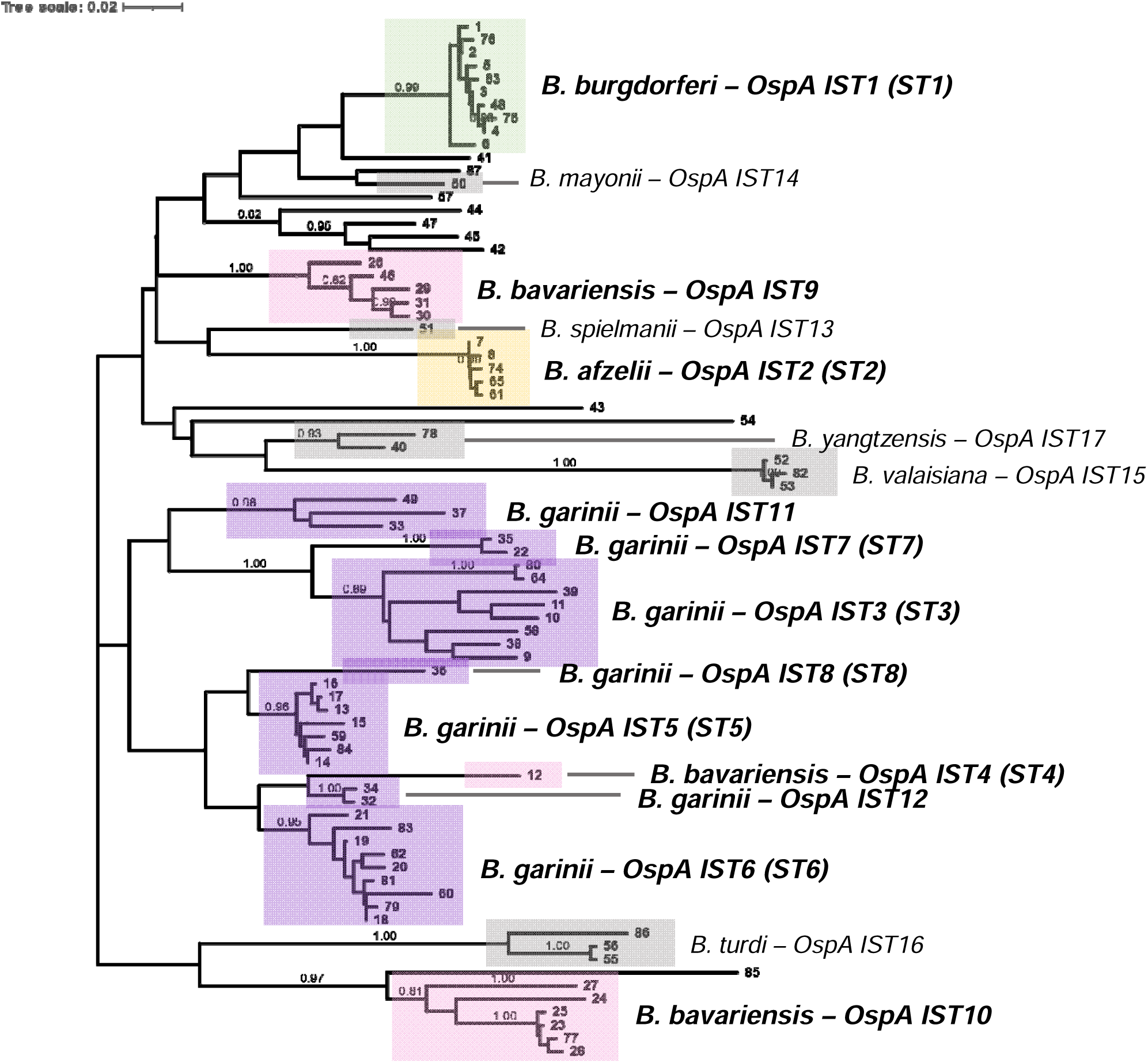
Phylogenetic tree of OspA variants. Phylogenetic analysis of 79 unique OspA protein variants. Clusters corresponding to the 17 ISTs defined in this study are labeled. The 12 OspA ISTs associated with species of *Borrelia* responsible for the majority of LD in North America, Europe, and Asia (*B. burgdorferi, B. garinii*, *B. bavariensis*, and *B. afzelii*) are bolded.

**Table 1.** Comparison of OspA *in silico* types (IST) and reported OspA serotypes corresponding to *B. burgdorferi* s.l. genospecies.

| ***In silico* Type (IST)** | **Serotype (ST)<sup>a</sup>** | **Genospecies** |
| --- | --- | --- |
| 1 | 1 | *B. burgdorferi* |
| 2 | 2 | *B. afzelii* |
| 3 | 3 | *B. garinii* |
| 4 | 4 | *B. bavariensis* |
| 5 | 5 | *B. garinii* |
| 6 | 6 | *B. garinii* |
| 7 | 7 | *B. garinii* |
| 8 | 8 | *B. garinii* |
| 9 | NR<sup>b</sup> | *B. bavariensis* |
| 10 | NR<sup>b</sup> | *B. bavariensis* |
| 11 | NR<sup>b</sup> | *B. garinii* |
| 12 | NR<sup>b</sup> | *B. garinii* |
| 13 | NR<sup>b</sup> | *B. spielmanii* |
| 14 | NR<sup>b</sup> | *B. mayonii* |
| 15 | NR<sup>b</sup> | *B. valaisiana* |
| 16 | NR<sup>b</sup> | *B. turdi* |
| 17 | NR<sup>b</sup> | *B. yangtzensis* |

<sup>a</sup> Serotypes listed here were classified using the monoclonal antibody-based typing system described by Wilske and colleagues (24, 25).
<sup>b</sup> NR = not reported

We next investigated the amino acid sequence diversity within and between ISTs. To begin, we measured the pairwise sequence identity across all 79 OspA variants. For each variant within an IST, we noted the highest sequence identity to variants from the same IST compared to the highest identity to variants not in that IST. The distributions of these two groups are shown in Figure 3 and Figure S1. In all 17 cases, a clear distinction was observed when comparing sequence identity within an IST to sequence identity of variants from another IST. This indicated that phylogenetic clustering was a robust method for assigning IST.

**Figure 3.**
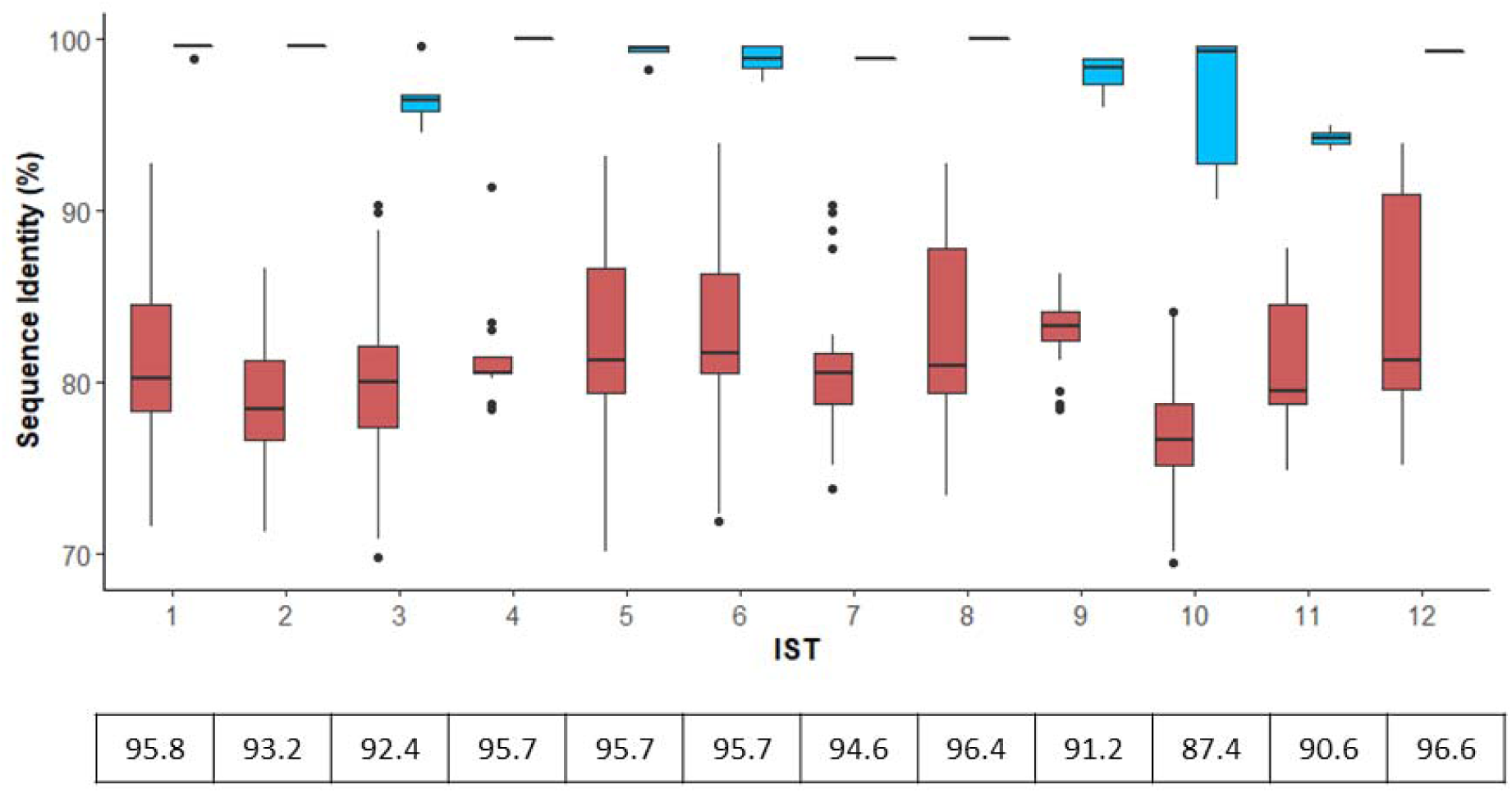
Comparison of OspA amino acid sequence identity within and outside of each IST. For each of the 79 OspA variants, sequence identity was compared to each variant from the same IST, and the highest percent similarity saved. The distribution of maximal sequence identity is shown in blue boxes. Likewise, sequence identity was compared to each variant not in the same IST and the highest percent similarity saved. The distribution of maximal sequence identity for these cases is shown in red boxes. A similarity threshold for each IST was then calculated as the midpoint between the tails of the two distributions, the values for which are depicted below the plot. As only a single OspA variant comprise both IST4 and IST8, sequence identity within these ISTs is depicted at 100%. Distributions for IST13-17 are found in Figure S1.

To further understand the variation between specific ISTs, we calculated the average sequence identity between OspA variants from each of the first 12 ISTs (Figure 4A) corresponding to *B. burgdorferi*, *B. afzelii*, *B. bavariensis*, and *B. garinii*. ISTs 4, 5, 6, 8 and 12 were found to have high between-IST sequence identity, reaching 93% average identity in the cases of IST6 and IST12. This was relatively unsurprising as these ISTs occupy the same branch in the OspA phylogeny (Figure 2). As variation in OspA is known to primarily occur at the C-terminal domain (20), we aligned these regions using consensus or representative OspA sequences these ISTs. We found that the C-terminal variable domain was relatively conserved across these ISTs, with the exception of a divergent segment from positions 206-230 (Figure 4B). This region of OspA contained 11 amino acid substitutions and one in/del, and clearly distinguished IST5 from IST8 and IST5 from IST6. In contrast, these substitutions were well conserved between IST4, IST6, and IST12. Similarly, we found that IST3 and IST7, which averaged 87.9% identity with one another, had well conserved C-terminal domains when consensus OspA sequences are compared to the other ISTs (Figure S2).

**Figure 4.**
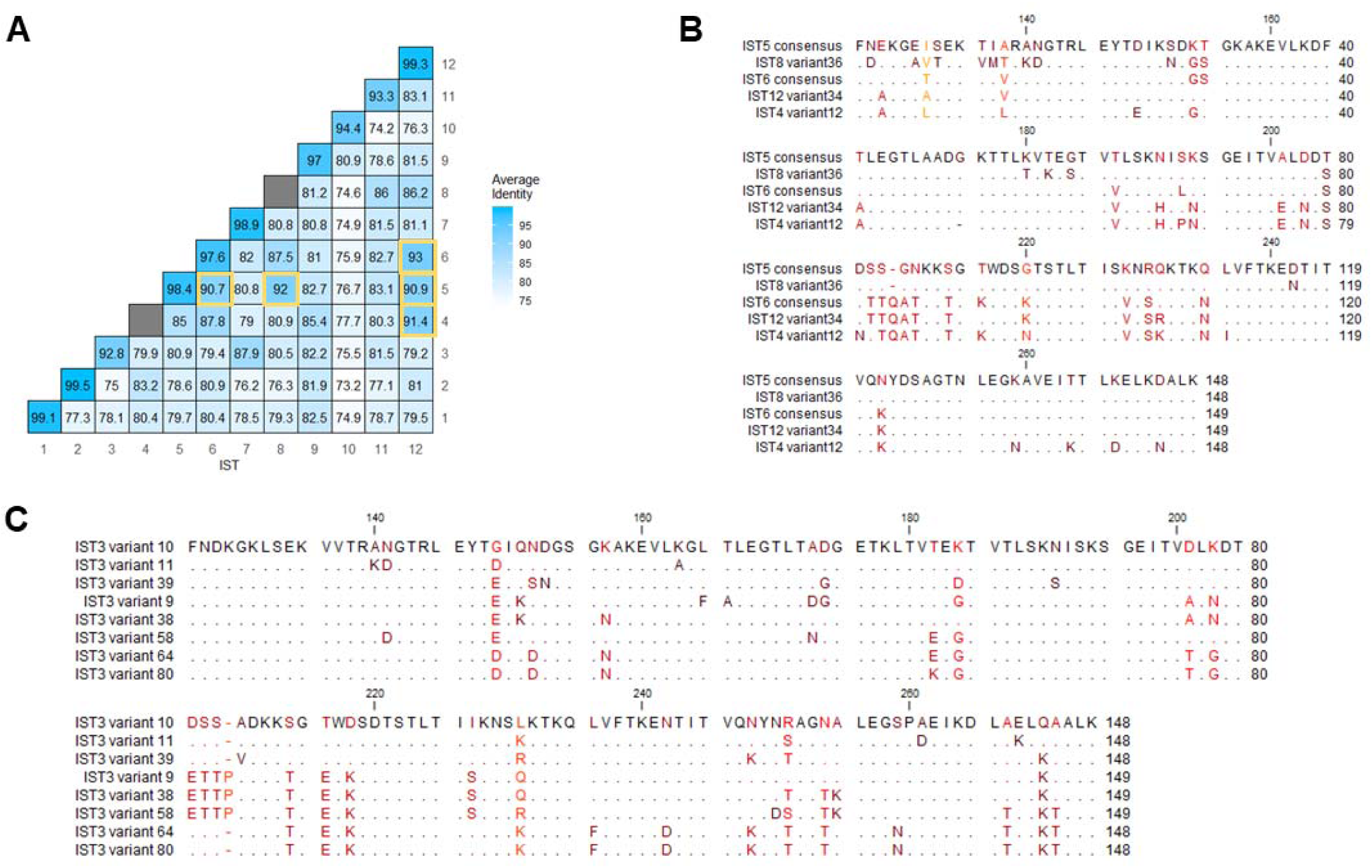
OspA sequence identity within and between ISTs. (A) Average percent sequence identity of OspA protein variants within and between IST1-12. As ISTs 4 and 8 consist of only a single OspA sequence variant, no within-IST identity was calculated for these types. (B) C-terminal domain alignment of ISTs 4, 5, 6, 8, and 12 OspA sequences. (C) C-terminal alignment of IST3 OspA variants.

Although within-IST sequence identity was consistently greater than between-ISTs, we found that IST3 OspA variants had the greatest sequence heterogeneity, only averaging 92.8% identity when compared with one another (Figure 4A). Based on the OspA phylogeny, it appeared that three sub-groups of IST3 were present in the collection (Figure 2) and were sufficiently diverged from one another to impact the average sequence identity. We aligned the C-terminal domain of the ten IST3 OspA variants and identified a region from positions 201-227 that characterizes the two sub-groups, consisting of eight AA changes and a single residue indel (Figure 4C). The full-length alignment of IST3 OspA variants is shown in Figure S3.

### Development of an *in silico* OspA typing pipeline

Leveraging this collection of diverse OspA variants, we sought to develop a standardized and automated method for determining the OspA IST of new isolates (Figure 5). To this end, we first tested if alignment of short-read NGS data of *ospA* from whole genome sequence could be used to generate a consensus sequence of OspA. For initial processing, a reference sequence(s) was needed for alignment. To determine if a single *ospA* sequence was sufficient as a reference for all ISTs, NGS reads of *ospA* from *B. burgdorferi* B31 (IST1) and a *B. garinii* IST6 isolate were simulated using SimuSCoP v1.0 and aligned to *ospA* sequences corresponding to ISTs 1-6. Coverage depth at each position of *ospA* is shown in Figure S4. While coverage was consistent when aligned to the *ospA* sequence of the respective ISTs, the same could not be said when aligning to heterologous reference sequences. Consistent with sequence divergence in the C-terminal region of OspA, alignment failed at the 3’ end of *ospA*. An example of this is illustrated in Figure S4A for *B. burgdorferi* when aligning to non-*burgdorferi ospA* sequences. Similarly, the *B. garinii* input reads were found to align poorly to the non-*garinii* reference sequences at the 3’ end (Figure S4B). Given that alignment quality is species- and IST-dependent, we concluded that alignment must be performed to corresponding reference *ospA* sequence from each target IST in order to maximize coverage and sequence integrity.

**Figure 5.**
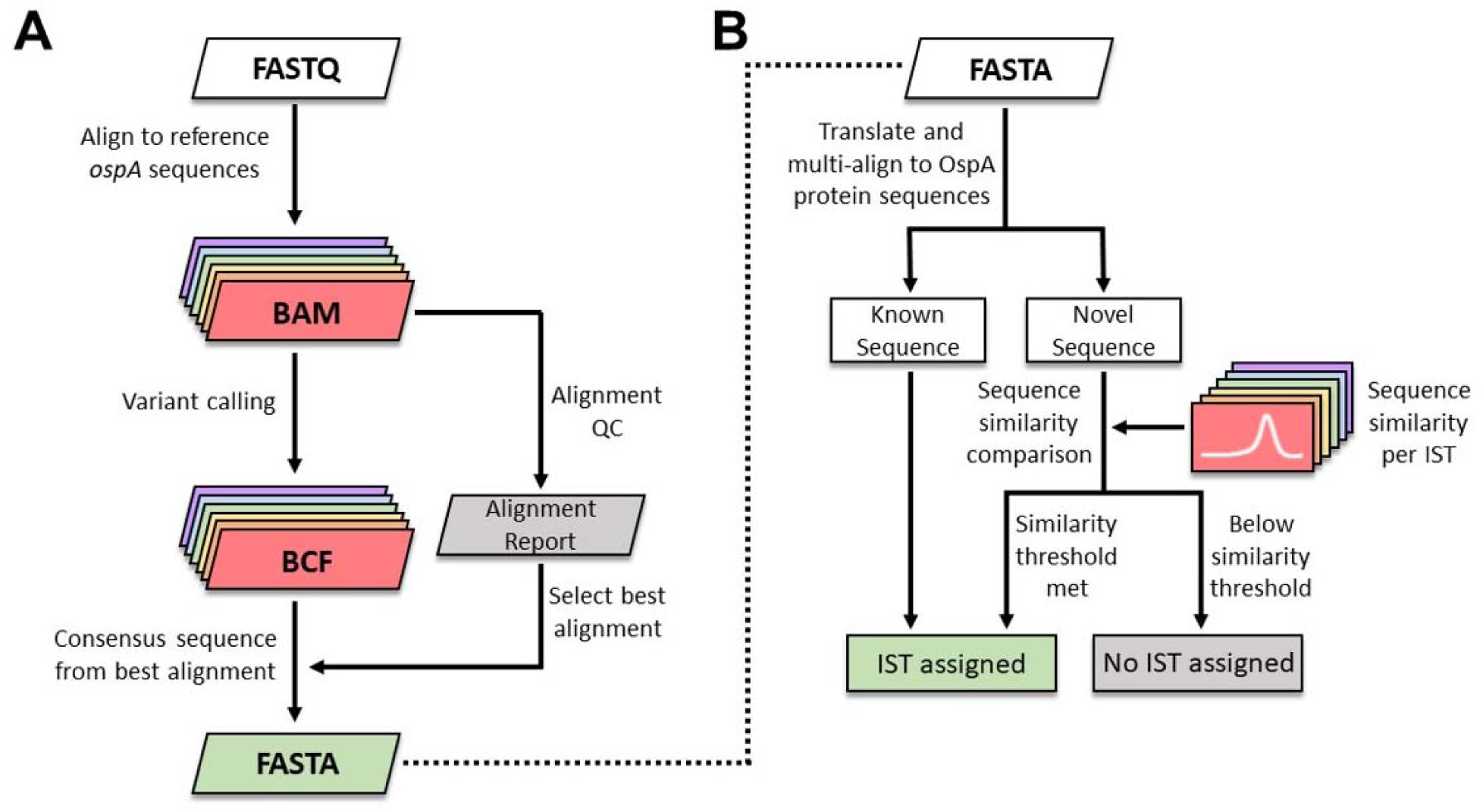
Workflow of an NGS-based *in silico* OspA typing pipeline. (A) NGS data of *ospA* are aligned to reference sequences for each IST. A consensus sequence is constructed based on the highest quality alignment. (B) The consensus *ospA* gene sequence is translated and multi-aligned against 79 known OspA variants. A sequence similarity threshold (Figure 3, Figure S1) is used to assign the IST of novel variants.

To determine the best alignment for consensus sequence building, both the total read coverage and coverage variation were considered. When aligning reads to a reference sequence of a different IST, the average coverage across *ospA* was found to be both lower and more variable, particularly in the C-terminal variable domain (Figure S3). To infer the best quality alignment across ISTs, an alignment score was calculated as the mean coverage minus one standard deviation. The alignment with the highest score was then used to build a consensus *ospA* sequence. Although this scoring selects for the most robust alignment, it did not consider cases where alignment quality or coverage is poor in general. In order to flag low confidence *ospA* alignments, minimum required values were set for total reads aligned, median read coverage, and depth of coverage across the entire *ospA* gene sequence (at least 5X sequence depth; see Materials and Methods). If the assembled genomic contigs are used as input, BLASTN was instead used to obtain alignments to the reference sequences of individual ISTs.

Finally, to determine the genospecies and IST, the generated consensus *ospA* sequence was translated and aligned to the collection of 79 OspA variants with species and IST assignments (Figure 5B). In cases where an exact OspA sequence match was available, the corresponding species and IST were assigned. If an exact match was not returned, the highest percent similarity to a known sequence was recorded alongside the IST associated with that sequence. As we had previously calculated the sequence identity distribution for OspA variants with known ISTs (Figure 3, Figure S1), these data were used to set thresholds for inclusion in each IST (see Materials and Methods). If the similarity of the query sequence was greater than or equal to the threshold for the known sequence, the associated IST was assigned. Otherwise, the sample was labeled as an Unknown IST.

### Testing of the OspA NGS pipeline using clinical *Borrelia* isolates

To evaluate performance of the OspA NGS pipeline, we ran a test dataset of whole genome sequences from 22 isolates representing ST1-6 (24) and six genospecies (*B. burgdorferi*, *B. garinii*, *B. afzelii*, *B. bavariensis, B. mayonii, and B. spielmanii*). A detailed comparison of the OspA ISTs identified by the pipeline is found in Table S4. The pipeline returned 100% concordance with the previously labeled genospecies, and all 22 of these isolates returned the expected results for OspA ST/IST (Table S4). A single *B. garinii* isolate, previously labeled as ST3, was found to harbor a novel allele of *ospA* with 99% identity to the reference IST3 *ospA* sequence used by the OspA NGS pipeline.

## DISCUSSION

A hexavalent (ST1-6) OspA-based vaccine, VLA15, is being developed for prevention of LD caused by these 6 most prevalent serotypes in North America and Europe (37). VLA15 is currently being tested for efficacy in a Phase 3 study conducted in both North America and Europe (NCT05477524). Determination of the OspA type of clinical isolates is important for vaccine development, both to determine concordance with the vaccine antigens, as well as to better understand the epidemiology and prevalence of different OspA types and *Borrelia* genospecies in human disease. Moreover, it is important to understand the genetic diversity of OspA. Characterization of the efficacy of VLA15 in Europe will be challenged by the number of potential OspA protein types present among European isolates. A reliable and implementable classification method that can readily accommodate new OspA types is currently lacking.

In the present study, we developed a bioinformatic pipeline that accurately assigns OspA ISTs and *Borrelia* genospecies from either OspA PCR amplicon sequence reads or assembled Borrelia genome data. These include IST1-IST8 which correspond to ST1-ST8 that had been previously assigned using traditional serological methods, as well as 9 novel ISTs (IST9-IST17). Our IST approach is rapid, simple, standardized, and only requires sequence data of *ospA* or *Borrelia* genome. The typing approach described in this work is the first reported pipeline specifically designed for OspA types of *B. burgdorferi* s.l.. Consistent with mAb-assigned OspA serotypes (24, 25), the OspA variants belonging to *B. burgdorferi* and *B. afzelii,* identified as IST1 and IST2, respectively, displayed near 100% within-group homology. New OspA clusters (IST11 and IST12) were identified for *B. garinii* as they diverge from the five established OspA serotypes associated with this species (25) (reported here as IST3 and IST5-IST8). We found the *B. bavariensis* IST4 European cluster was geographically distinct from two novel clusters (IST9-IST10) containing *B. bavariensis* isolates from Asia. The divergence between these OspA groups is expected given evidence for a clonal expansion of the European population of *B. bavariensis* (38–40). Only a single OspA variant corresponding to IST4 (*B. bavariensis*) and IST8 (*B. garinii*) was identified in our sequence collection, and thus, within-serotype similarity was set at 100% for these ISTs.

Traditional *Borrelia* serotyping by OspA-specific mAbs enabled the identification of ST1-8 (25) and serotypes J1 to J11 among Japanese *B. burgdorferi* s.l. isolates (41). The utility of mAb-based typing is technically challenging and subject to the level of expression of the OspA protein *in vitro* (24) which can affect mAb binding. In addition, unlike other bacterial species (*e.g.,* streptococci), standardized mAb typing reagents are not available. Moreover, some of the OspA-specific mAb reagents have been prone to cross-reactivity due to high sequence homology. As an example, cross-reactivity between *B. burgdorferi* and *B. bissettii* has been reported using H3TS, an OspA-specific mAb used for the identification of OspA ST1 (42). By comparison, the IST pipeline described herein unambiguously differentiates *B. burgdorferi* as IST1 based on OspA sequence. Since the IST OspA pipeline only scans for *ospA* sequences, interference of host DNA or DNA of similar pathogens lacking *ospA*, such as relapsing-fever *Borrelia,* is also minimized. Consequently, the pipeline is capable of IST assignment using a small amount of *Borrelia* DNA in a background of host DNA. Only NGS reads aligned to *ospA* are utilized, and IST assigned with high sensitivity and specificity.

Genotyping *Borrelia* isolates in clinical samples is desirable for vaccine development to better understand the epidemiology of LD. *Borrelia* genospecies and OspA types are differentially distributed across the northern hemisphere and have been associated with distinct manifestations of LD (24, 25, 43, 44). Sequence alignment of the *ospA* gene and analyses based on OspA amino acid sequence similarity have been used broadly for clustering *Borrelia* isolates (22, 41, 45–47). This clustering has provided useful genospecies information in agreement with classification based on the analysis of other conserved chromosomal genes (27) and mAb-based serotyping (24). However, alignment analyses suffer from limitations of scalability and standardization, and often partial *ospA* gene was targeted. The pipeline presented here provides high resolution to differentiate between multiple OspA ISTs, as well as the ability to identify novel phylogenetic IST clusters within a single genospecies (*e.g.,* IST9 and IST10 for *B. bavariensis* and IST14 and IST12 for *B. garinii*). The ability for *de novo* strain detection, identification, and assignment of *ospA* alleles is an advantage of NGS-based typing platform, with the capability to easily build on phylogenetic trees as new disease-causing *Borrelia* species are discovered (48) and more complete genomes are sequenced.

Evaluation of the OspA IST pipeline was limited to 22 *Borrelia* isolates from a single collection. The lack of *Borrelia* isolates with both serological typing data and OspA sequence information presented a challenge. In addition, the current pipeline design does not take co-infection with multiple *Borrelia* strains into consideration, and therefore can detect the dominant IST only when working on samples that may carry >1 isolates from different ISTs. As this pipeline relies on the association between OspA sequence and genospecies, assignment of genospecies based on IST could potentially be occluded in the case of horizontal gene or plasmid transfer (49).

Our work has uncovered novel clusters of OspA variants among strains of *B. burgdorferi* s.l., and also introduced the concept of ISTs as a new nomenclature for strain characterization. Furthermore, we developed an open-source reliable OspA analysis pipeline which enables characterization of novel *Borrelia* OspA types using NGS data without the need for traditional, antibody-based serotyping systems. As global surveillance of *Borrelia* continues to expand, this method has the potential to document additional ISTs. Such novel ISTs will be included in future updates to the *in-silico* OspA typing pipeline.

## AUTHORS’ CONTRIBUTIONS

All authors met ICMJE criteria for authorship and participated in the study design and conceptualization (JTL, PAL, LH, RS, ASA, JWH), methods development (JTL, LH, BSP, DK), data collection and interpretation (JTL, LH, ZL, LDN, VR, BSP, DK, LA, KEL, UR), writing the original draft (JTL, LH, ZL, JWH, RS, PAL), and study management (LH, RS, PAL, ASA). All authors contributed to the development of the manuscript.

## DATA AVAILABILITY

The internal datasets generated and/or analyzed during the present study are available in the NCBI repository BioProjects: PRJNA1041728. Accession numbers and isolate names are included in Table S1. Code for the *in silico* OspA pipeline will be available on GitHub ( https://github.com/pfizer-opensource/) following acceptance of the manuscript.

## ACKNOWLEDGEMENTS

The authors thank University of Amsterdam colleagues, Alje van Dam, for his role in the isolation of Dutch *Borrelia* spirochetes from human samples, and Amber Vrijlandt, for the stock preparation and sorting of all Dutch *Borrelia* isolates. The authors also acknowledge Christina D’Arco (Pfizer) for writing support, and colleagues at Valneva (Urban Lundberg and Andreas Meinke) as well as Gary Wormser (New York Medical College) for providing isolates.

## Supplementary Data

**Table S1. Sample and NGS metadata of internally sequenced *Borrelia* isolates.**

**Table S2. Sample and NGS metadata of publicly obtained *Borrelia* isolates.**

**Table S3. OspA variants for IST pipeline.**

**Table S4. Evaluation of the IST pipeline against isolates with known ST and genospecies.**

**Figure S1.**
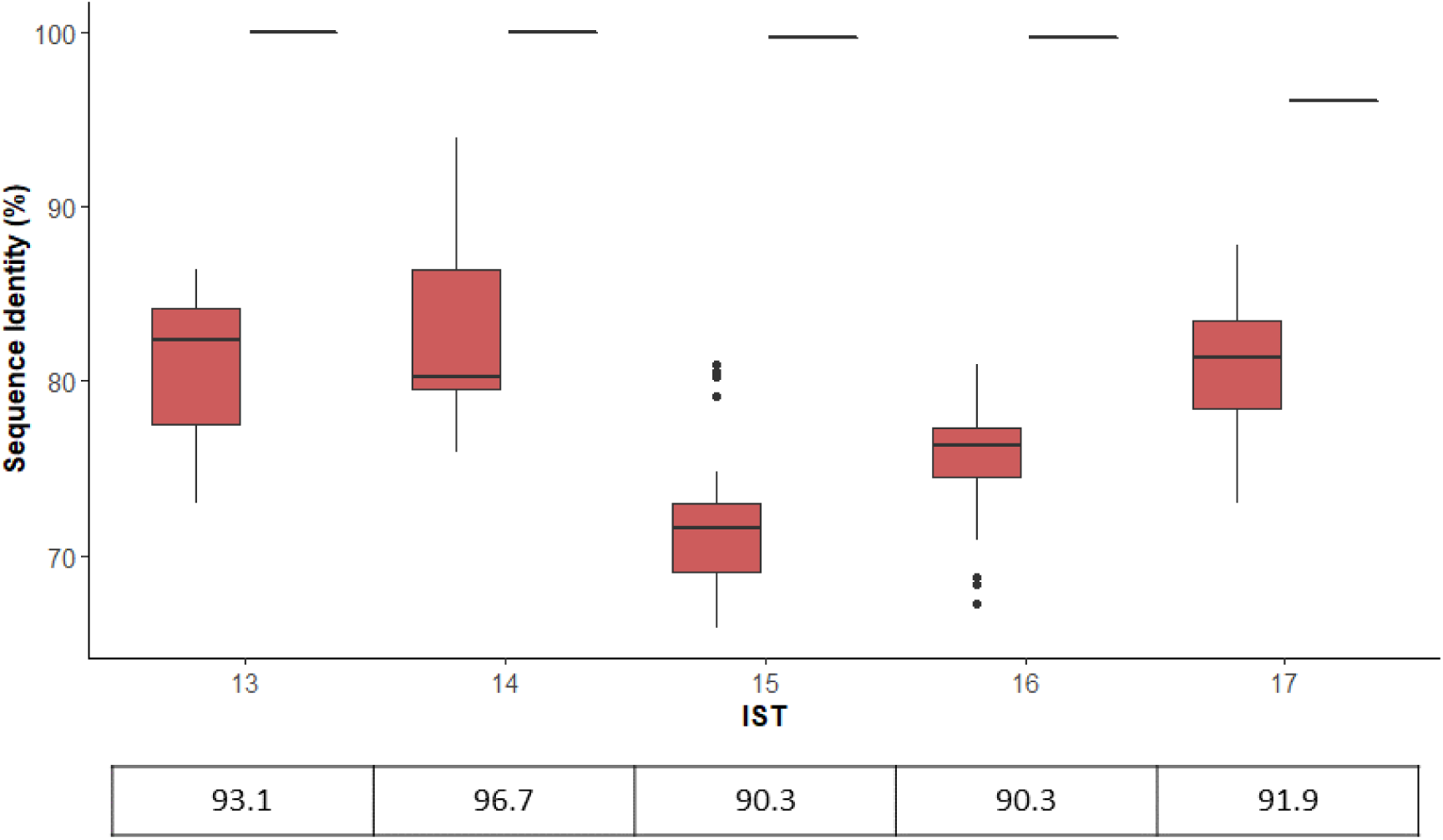
Comparison of OspA sequence identity for ISTs 13-17.

**Figure S2.**
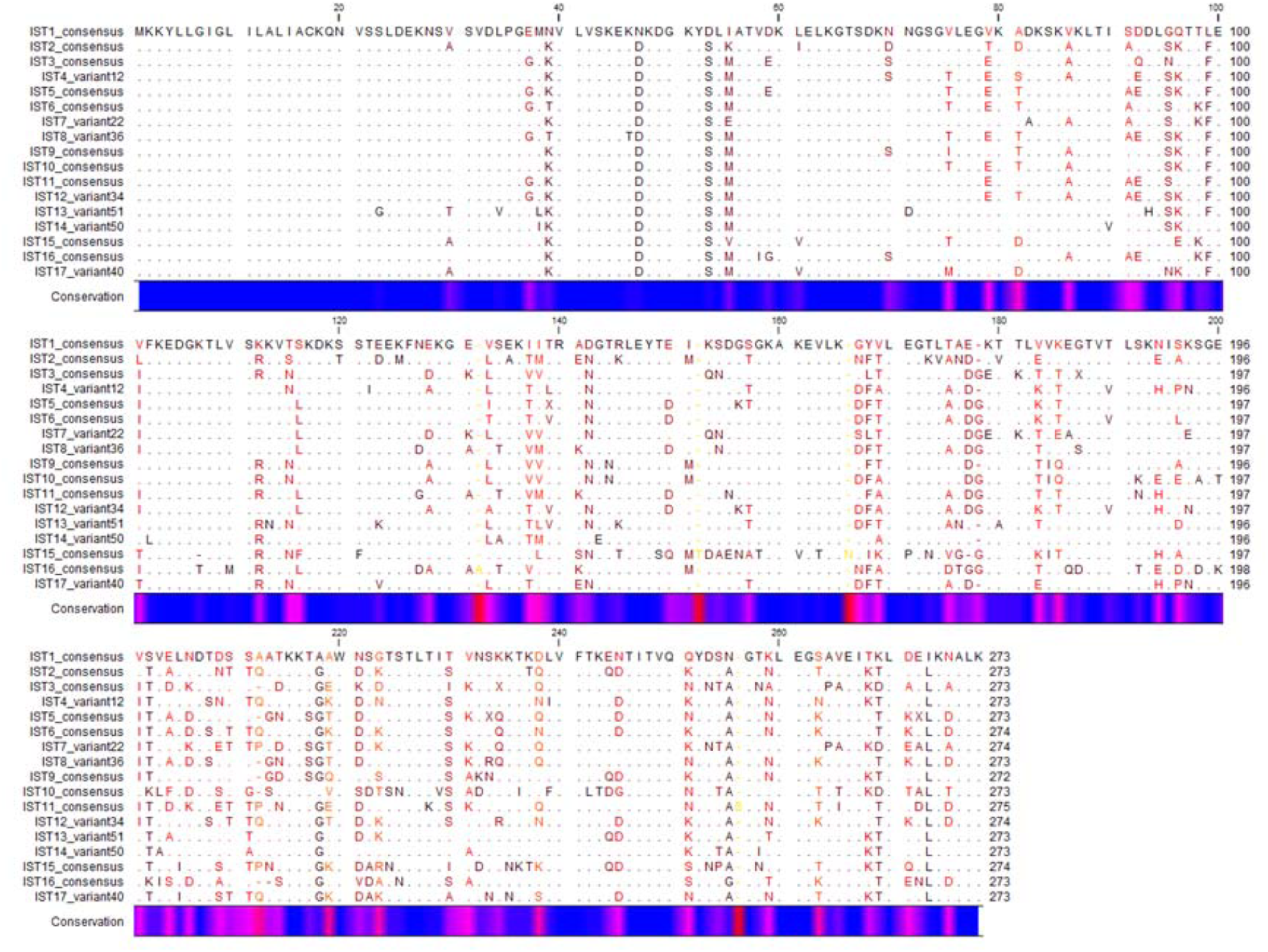
Alignment of IST consensus OspA sequences.

**Figure S3.**
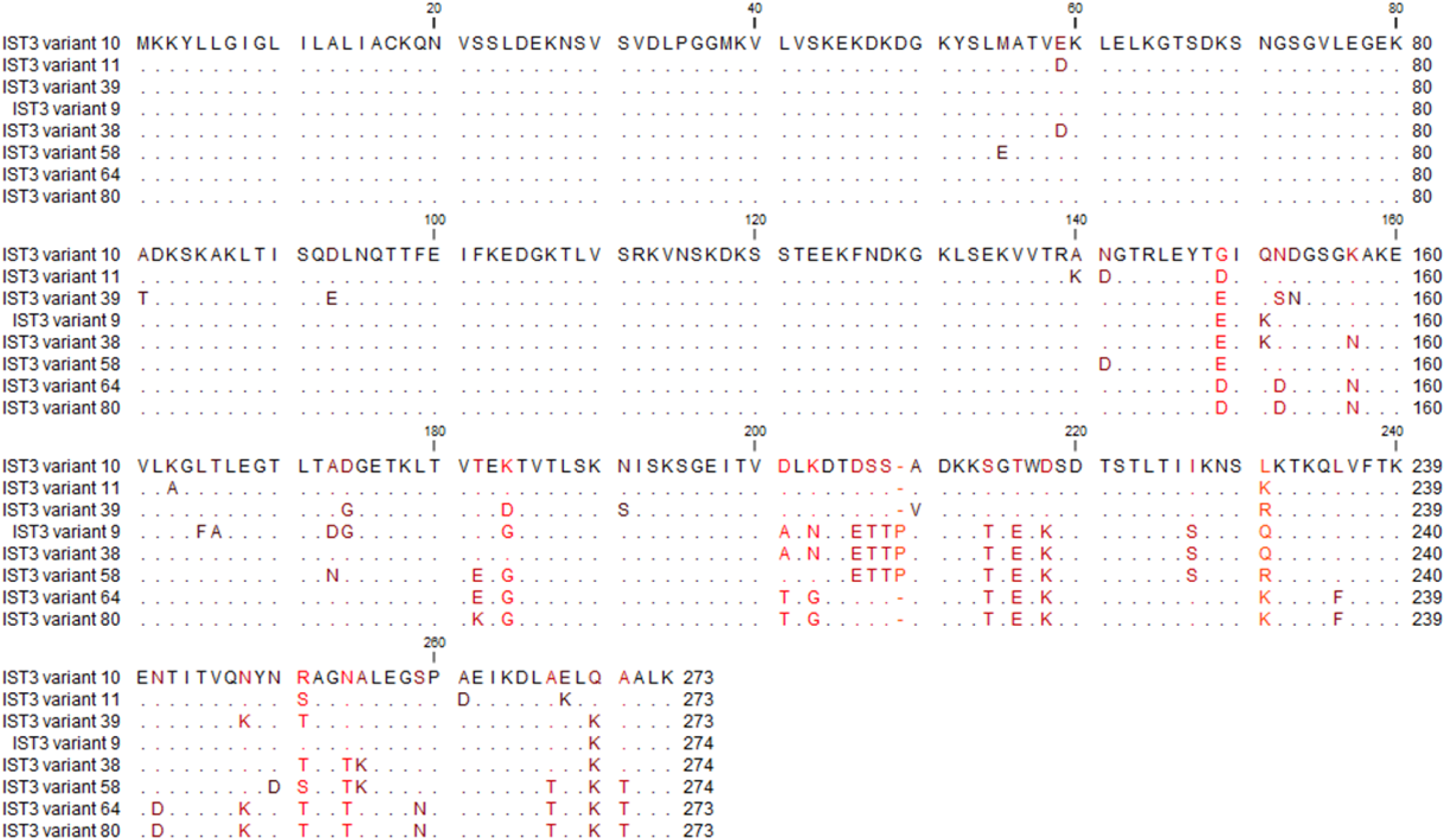
Alignment of IST3 OspA sequences.

**Figure S4.**
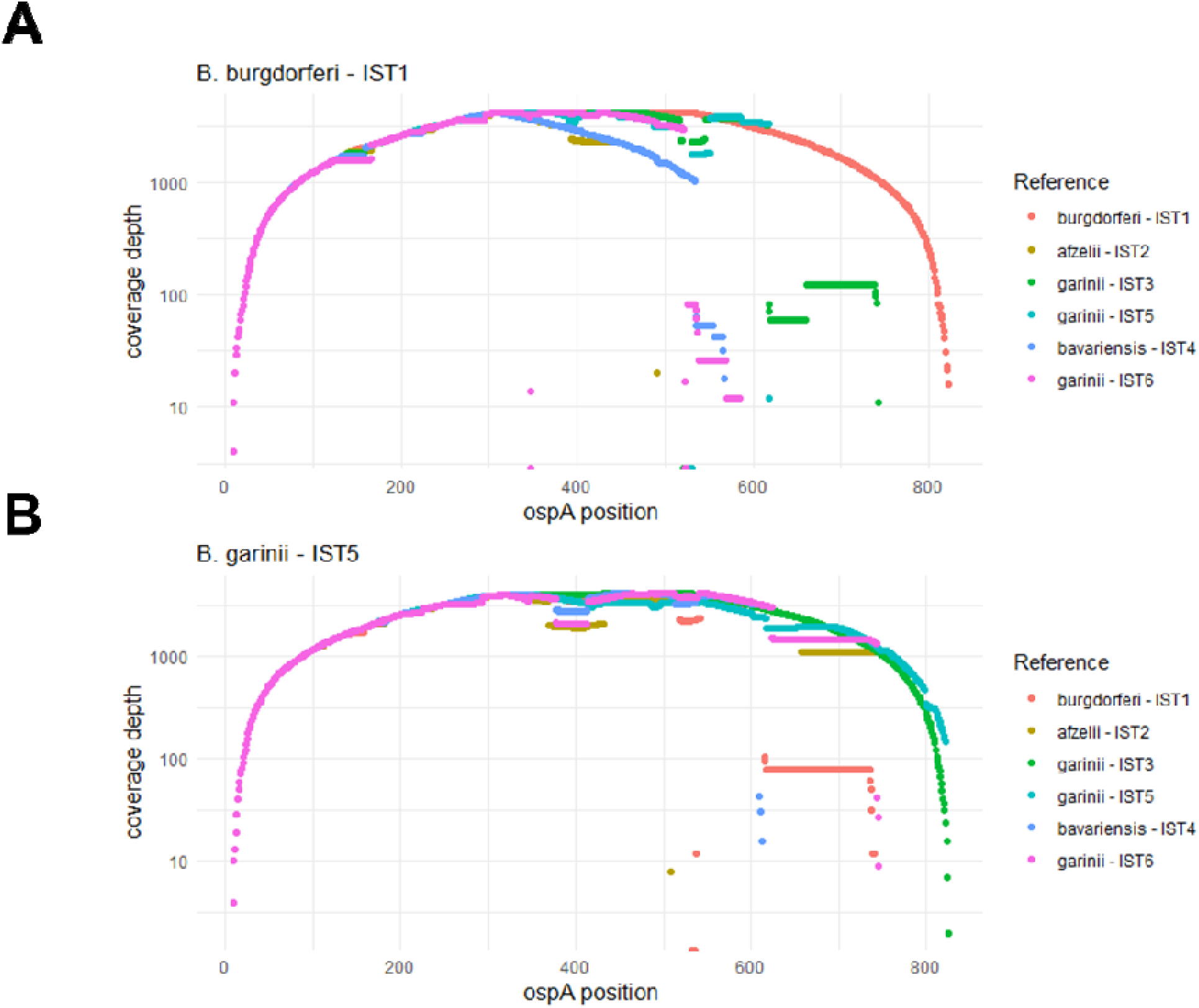
Simulated NGS coverage for a *B. burgdorferi* and *B. garinii* isolate.

